# Traveling Wave Analysis of a Go-or-Grow Invasion Model with ECM-Regulated Phenotypic Switching

**DOI:** 10.64898/2026.04.23.720361

**Authors:** Gopinath Sadhu, Mohit Kumar Jolly, Philip K Maini

## Abstract

Experimental studies show that tumor cells adopt migratory or proliferative phenotypes depending on the local extracellular matrix (ECM). In this work, we propose a minimal go-or-grow invasion model, comprising two specialist cell phenotypes: proliferating and migratory, with phenotypic switching and cell migration depending on local ECM density. Numerical simulations of this model reveal that the spatial arrangement of proliferative and migratory cells depends on the choice of phenotypic switching function. We then ask whether this specialist cell-phenotype model can be reduced to a generalist cell-phenotype model. We derive a relationship between the reduced model and go-or-grow model in the fast phenotypic switching regime. We observe that the reduced model captures the dynamics of the original model, for a range of realistic phenotypic switching functions. We analytically derive the minimum traveling wave speed of the reduced model in a homogeneous ECM bed. Moreover, using linear stability analysis on the go-or-grow model, we recover the same wave speed expression. In addition, we numerically explore how the key parameters influence the traveling wave speed profile. Our analysis indicated the counter-intuitive result that the wave speed is independent of the matrix degradation rate, and our simulations show that, at most, the speed is weakly dependent on this parameter.

## 1. Introduction

Tumors consist of diverse populations of cells that exhibit various phenotypes, including migratory and proliferative cells. These cells can switch phenotypes in response to distinct driving forces in the tumor microenvironment. For example, hypoxia, a shortage of oxygen, occurs due to high oxygen consumption at the tumor periphery and is a factor in cells switching their phenotype during tumor invasion [1]. Numerous studies have shown that hypoxia regulates the epithelial-to-mesenchymal transition (EMT), in which epithelial cells, which are typically less migratory and more proliferative, transition to a mesenchymal phenotype characterized by increased migratory potential and reduced proliferation [2, 3, 4, 5]. Additionally, hypoxia influences metabolic processes, prompting tumor cells to shift their glucose metabolism from oxidative phosphorylation to glycolysis. This switch results in excessive lactate, H+ ions, and CO_2_ production, leading to an acidic tumor microenvironment. The acidity of this microenvironment also plays a critical role in phenotypic switching [6]. Furthermore, experimental studies showed that when cells are in a highly crowded cell area, they may inhibit proliferation but activate the migration mode in response to nutrient limitation [7, 8]. Recently, experimental studies have shown that the extracellular matrix (ECM) also controls phenotypic switching between cell types [9, 10]. However, the mechanisms underlying how these phenotypic switches influence tumor migration dynamics remain poorly understood.

Numerous mathematical models have been proposed to elucidate the dynamics of tumor growth and invasion [11, 12, 13]. Among those models, the go-or-grow modeling framework, where one cell population only proliferates, and another cell population only migrates, is the most popular phenotypic model to study cancer invasion [14, 15, 16]. El-Hachem et al. [17] proposed an invasion model in which a single species invades the surrounding ECM medium, and cell diffusion is influenced by the ECM’s volumetric effects. This modeling framework is extended in the go-or-grow architecture by incorporating heterogeneity in tumor cell phenotypes [12], and it is assumed that cellular migration is hindered by the collective volumetric effects of local cells and ECM densities. In addition, phenotypic switching is based on local tumor microenvironmental factors, such as total cell density, spatial constraints, and ECM density [8, 18]. Recently, Falcó et al. [8] analyzed the traveling-wave behavior of a go-or-grow model in which phenotypic switching between the populations depends on the total cell density. Crossley et al. [18] proposed a phenotypically structured invasion model of tumor invasion in which phenotypic switching depends on ECM density. However, how ECM-dependent phenotypic switching affects invasion properties (such as speed) in a go-or-grow model remains poorly understood.

In this work, we propose a go-or-grow model of tumor invasion into an ECM bed in which the local ECM density regulates phenotypic switching. The model consists of a coupled system of three partial differential equations (PDEs) describing two specialist cell types: proliferating cells and migratory cells, and the ECM. We assume that the rates of phenotypic switching between proliferative and migratory states can either be constant, or depend on local ECM density: tumor cells switch from a proliferative to a migratory state when ECM density increases, while migratory cells revert to a proliferative phenotype when ECM density decreases [19, 20]. The diffusion of migratory cells depends on the local ECM density. This study investigates the conditions under which the go-or-grow model for two specialist cell phenotypes can be reduced to a single-phenotype cell model, and examines the relationship between the two models. We confirm that the behavior of the go-or-grow model is accurately captured by the reduced model in the fast-phenotype-switching regime by numerical simulations across different choices of phenotypic switching functions. We also find that in this regime, linear analysis of the traveling wave far ahead of the front yields an expression for the minimum wave speed identical to that derived for the reduced model. However, this analysis also reveals a positive eigenvalue suggesting that, counter to standard traveling wave analysis, the steady state far ahead of the wave is unstable in the traveling wave coordinate system. We hypothesize that the behavior of the solution far behind the wavefront forces it to lie in the stable manifold of the solution far ahead of the front. This is confirmed by linear analysis and numerical simulations of the go-or-grow model.

## 2. A go-or-grow invasion model

Let *m*(**x**, *t*), *p*(**x**, *t*) and *e*(**x**, *t*) denote the densities of migratory cells, proliferative cells and ECM, respectively, at time *t* and position **x** = (*x, y, z*) in cartesian coordinate space and define the total cell density by

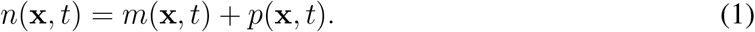

A minimal go-or-grow invasion model, where phenotypic switching and proliferation may depend on local ECM density, and ECM is degraded by migratory cells, is given by

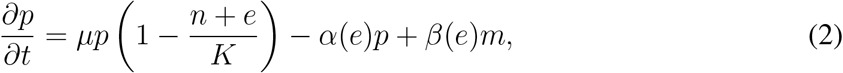

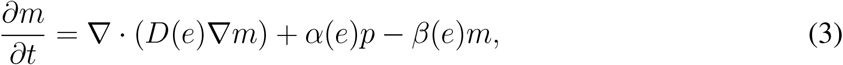

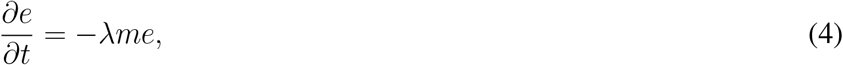

where 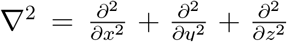. The proliferating cells proliferate at a linear rate *µ* logistically, depending on the ECM and total cell densities with carrying capacity *K*. The proliferative cells switch into migratory cells either at a constant rate, or as an increasing function *α*(*e*) of ECM density [19]. Phenotypic switching from migratory to proliferative cells is either constant or a decreasing function *β*(*e*) of ECM density. Experimental studies reported that the surrounding ECM hinders tumor migration [21], so, we take the diffusivity of migratory cells, *D*(*e*), to be a decreasing function with *D*(*e*) *<* 0. The last equation models ECM degradation due to migratory cells with a degradation rate *λ*.

For the rest of the paper, we will refer to this model as the go-or-grow model.

### 2.1. Model reduction to a mixed cellular phenotype model for invasion

Experimental studies reported that the time-scale of switching between proliferative and migratory phenotypes can be faster than that of cellular-level dynamics, such as cell proliferation, cell migration, and ECM degradation [22]. We define the parameter *ε* to be the ratio between the phenotypic switching time scale and the population dynamics time scale, namely

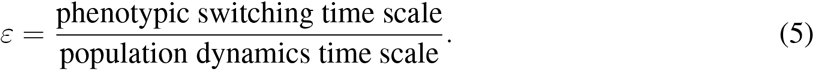

The condition 0 *< ε* ≪ 1 implies that phenotypic adaptation occurs much faster than migration or proliferation. Fast phenotypic switching is modeled for Eqs. (2)-(3) by assuming [8, 23]

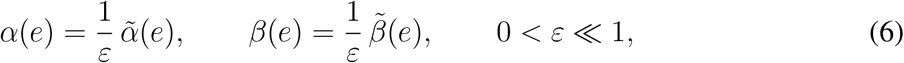

where 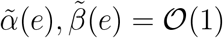.

Hence, the typical fast-switching system takes the form

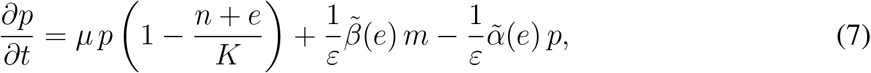

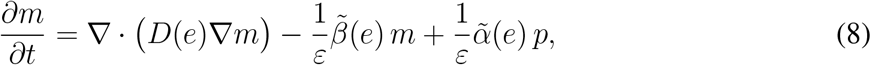

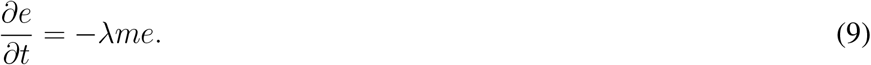

In the limit *ε* → 0, the leading-order balance yields the relation

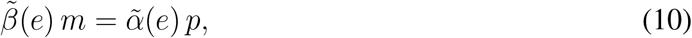

and the two cell densities *p* and *m* can be related to the total cell density (*n*) via

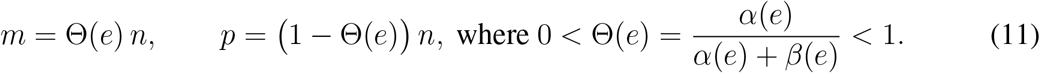

Substituting these expressions into the total density equation (sum of Eqs. (7) and (8)) leads to the reduced invasion model

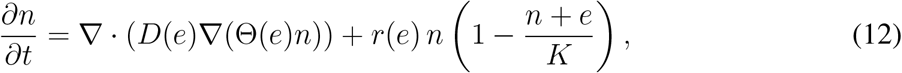

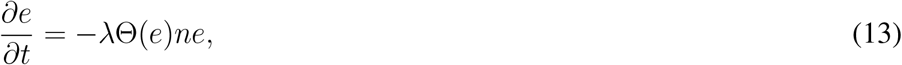

where

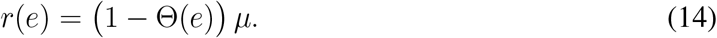

In the rest of the paper, we refer to this model as the reduced model.

When *λ* = 0, the ECM will have a fixed density, i.e., *e*(*x, t*) = *e*_0_. Then Eq.(12) becomes

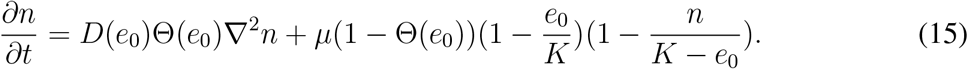

This has the form of the classic Fisher-KPP equation.

## 3. Numerical results

In this section we demonstrate, through taking specific forms of phenotypic switching functions, that the reduced generalized-phenotype model can capture the go-or-grow invasion model, in the fast phenotypic switching regime. To solve the model (Eqs. (2)-(4)) and reduced model (Eqs. (12)- (13)) we consider the one-dimensional case with domain [0, *L*] along the positive x-axis. To obtain numerical results, we employed the forward time central difference in space (FTCS) scheme within a finite difference framework in a one-dimensional Cartesian domain [0, *L*] (details are described in Appendix A). For the specialist phenotype model (Eqs. (2) and (3)), we assumed that the tumor initially grows solely from proliferative cells. Hence, we take initial conditions *p*(*x*, 0) = 0.5 for 0 ≤ *x <* 2 and *p*(*x*, 0) = 0 for *x >* 2, *m*(*x*, 0) = 0 and *e*(*x*, 0) = *e*_0_ for *x* ∈ [0, *L*] with *L* = 10. For the migratory cells (*m*(*x, t*)), we employ zero-flux conditions at the boundary points *x* = 0 and *L*. For the mixed phenotype model (Eq. 12), we use zero-flux conditions at the boundary points [0, *L*] and initial condition *n*(*x*, 0) = 0.5 for 0 ≤ *x <* 2 and *n*(*x*, 0) = 0 for *x >* 2, and *e*(*x*, 0) = *e*_0_ for *x* ∈ [0, *L*].

Figure 1 displays the spatial distribution of specialist phenotype cells: proliferative (*p*) and migratory cell (*m*), generalized phenotype cell (*n*), and ECM (*e*) densities. It is observed that our reduced model captures the behavior of the go-or-grow model for each choice of phenotypic-switching functional form. In addition, it is interesting to observe that proliferative and migratory cells can coexist throughout the tumor when phenotypic switching is constant, or switching from migratory to proliferative occurs only in response to local ECM density (Figures 1i and iv). When the phenotypic switching function that drives proliferative cells to become migratory depends on ECM density, migratory cells are at the tumor periphery (Figures 1ii, iii and v). In addition, we tracked the position of the tumor front across different functional forms of phenotypic switching (Figure 1vi). We observe that tumor invasion becomes more aggressive when the transition rate from proliferative to migratory cells is independent of the ECM. In contrast, the reverse transition (migratory to proliferative) remains ECM-dependent. In contrast, when the transition from migratory to proliferative cells is independent of ECM density, and the proliferative-to-migratory switch depends on the ECM, tumor invasion is slowed.

**Figure 1:**
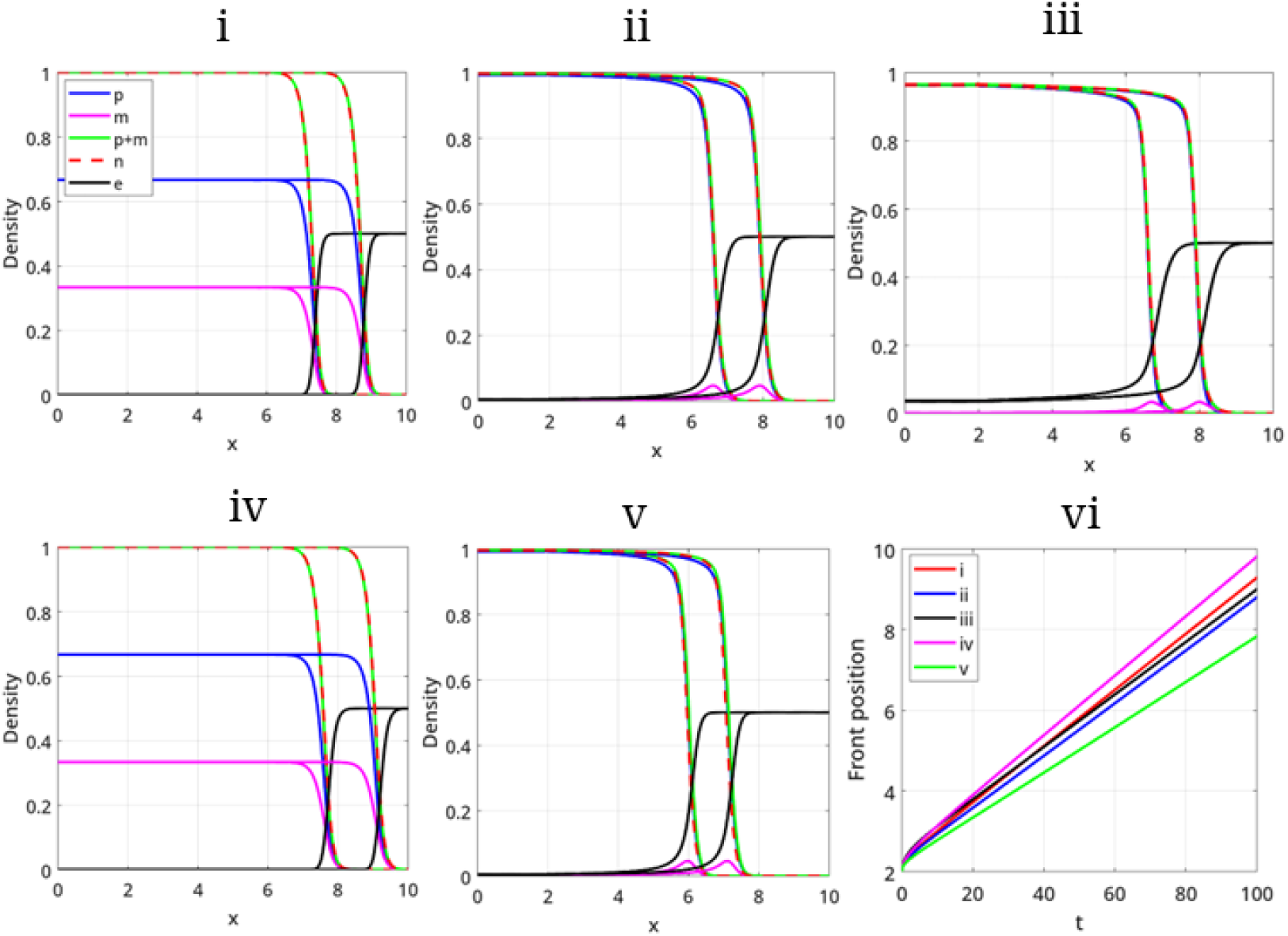
Numerical solutions for the two models given by equations Eqs. (2)-(4) and Eqs. (12)-(13). The spatial distribution of *p, m, p* + *m* and *n* at t = 40 and 80 for (i) *α*(*e*) = 10, *β*(*e*) = 20, (ii) *α*(*e*) = 10*e, β*(*e*) = 20(1 − *e*), (iii) 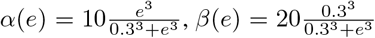, (iv) *α*(*e*) = 10, *β*(*e*) = 20(1 − *e*) and (v) *α*(*e*) = 10*e, β*(*e*) = 20. Other parameter values are *e*_0_ = 0.5, *µ* = 1 and *λ* = 5, *K* = 1 and 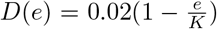. (vi) Comparison of tumor front point propagation over time for different choices of phenotypic switching functions.

## 4. Traveling wave analysis for specialist phenotype cell populations

We now consider the go-or-grow model equations (Eqs. (2)-(4)) in a one-dimensional Cartesian coordinate system to analyze constant profile, constant speed, traveling wave solutions of the form: *p*(*x, t*) = *U*_1_(*z*), *m*(*x, t*) = *U*_2_(*z*) and *e*(*x, t*) = *U*_3_(*z*), where *z* = *x* − *ct* ∈ (−∞, ∞) and *c >* 0 is the constant wave speed. Hence, the system Eqs. (2)-(4) in the traveling wave coordinate becomes

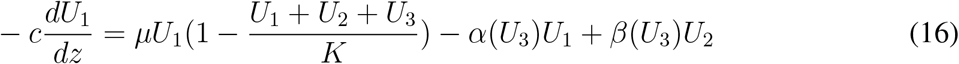

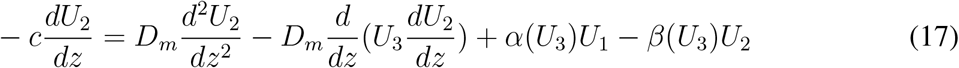

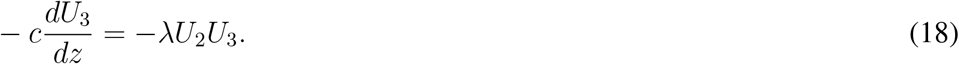

The steady state equations for the above system are

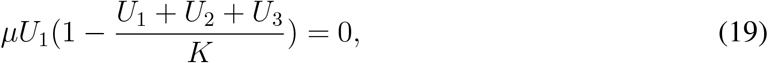

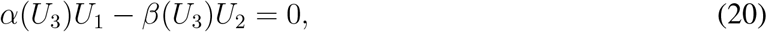

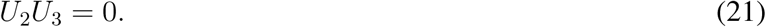

The solutions to these are (*U*_1_, *U*_2_, *U*_3_):

SS1: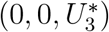, where 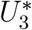 is an arbitrary constant.

SS2: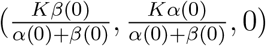.

SS3: 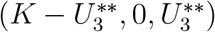 with 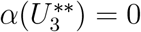.

Note, SS1 is precisely the state we observe far ahead of the wave, with 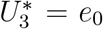, while SS2 and SS3 are the states we observe behind the front. It is interesting to notice that SS2 corresponds to panel (i) and (iv) of Figure 1. Clearly, for the choice of linear (*α* = 10*e*) or Hill 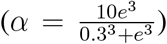 types, 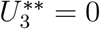, hence the steady state (SS3) is (*K*, 0, 0), which is observed in Figure 1 ii and v. It is observed from Figure 1(iii) that the system has not yet reached a steady state. We have shown on a larger domain that the ECM in the tumor region does indeed tend to zero over time (SI Figure S1).

### 4.1. Dispersion relation

Far-ahead of wave, the steady state is *U*_1_ = *U*_2_ = 0 and *U*_3_ = *e*_0_. For linear stability analysis, we consider *U*_1_ = 0+*Ũ*_1_, *U*_2_ = 0+*Ũ*_2_ and *U*_3_ = *e*_0_+*Ũ*_3_ where |*Ũ*_*i*_| ≪ 1 for *i* = 1, 2, 3. Substituting into Eqs. (16)-(18) and ignoring higher order terms, on dropping the’~‘, the linearized system takes the form

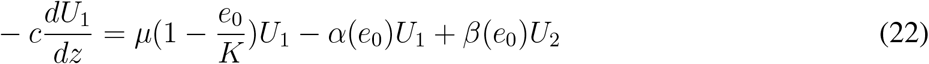

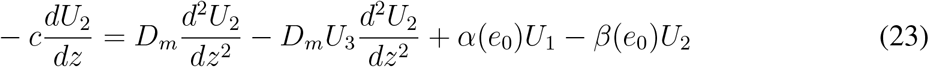

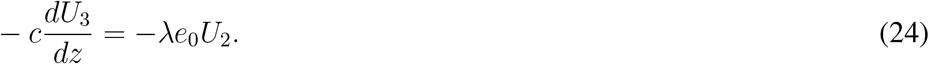

By substituting *U*_1_(*z*) ∼ *A* exp(*σz*), *U*_2_(*z*) ∼ *B* exp(*σz*) and *U*_3_(*z*) ∼ *D* exp(*σz*), where *A, B* and *D* are constant, we obtain

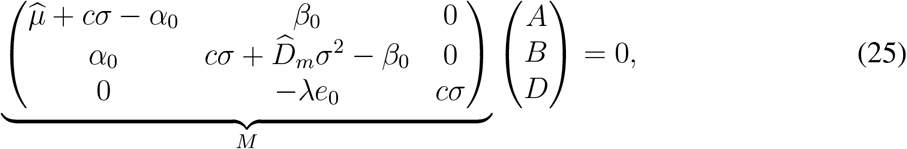

where 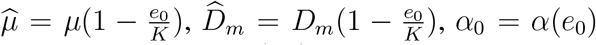 and *β*_0_ = *β*(*e*_0_). In order to ensure that *A, B, D* ≠ 0, we require det(*M*) = 0, which gives

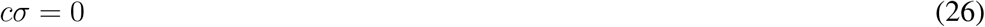

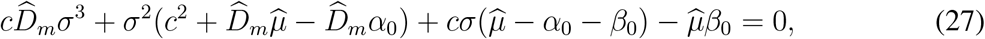

We now make the following observations:

1. *σ* = 0 implies SS1 can, at best, be neutrally stable.
2. From Eq. (27), it is clear that one root is positive, making SS1 unstable, implying that, contrary to our simulation results, we should not be observing a constant profile, constant speed, traveling wave.
3. We require the remaining two roots to be real and negative (employing the usual non-negativity of solutions argument).
4. The stability of the steady state, and hence the wave speed, is independent of *λ*.

Standard analysis of a cubic (see Appendix B) shows that we need 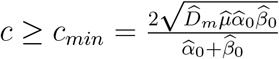. for the two other roots of *σ* to be purely real.

Note that Eq. (27) is independent of *λ* implying that the go-or-grow model will have a wave speed *c* that does not depend on *λ*.

Hence, we are left with the issue of the root that is positive, implying an instability. While the system is too complicated to analyze in general, we consider the fast phenotypic regime, 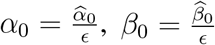, where 0 *< ϵ* ≪ 1 and 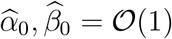, Eq. (27) becomes

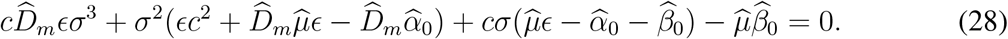

As *ϵ* → 0, this reduces to

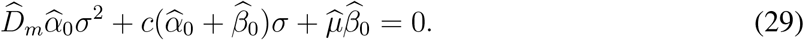

This equation has solutions with negative real part, but for purely real solutions, we require the discriminant to be non-negative, hence we derive the condition 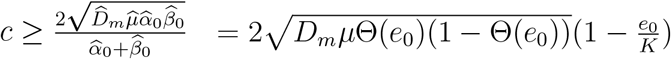. This same expression can be obtained in an alternative way, as shown in Appendix B and, in fact, also arises from Eq. (15), which is the classical Fisher KPP equation. Indeed for the reduced model, far-ahead of the wave, even with a non-zero value of the matrix degradation rate *λ*, the linear analysis still predicts a wave speed that is independent of *λ*.

Under this condition on *c*, Eq. (27) has one positive real root and two negative real roots; *σ*_1_, *σ*_2_ *<* 0 and *σ*_3_ *>* 0. So, *σ*_1_, *σ*_2_ and *σ*_3_ are eigenvalues, and we denote their corresponding eigenvectors by *v*_1_, *v*_2_ and *v*_3_, respectively. Hence

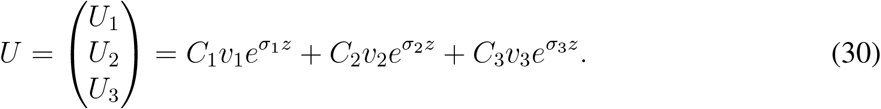

In the fast phenotype regime, the eigenvectors for *σ*_1_ and *σ*_2_ are 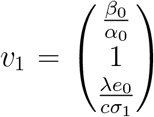 and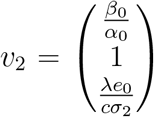, respectively.

Note that far behind the wave 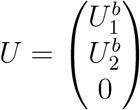 where 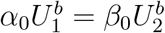. Hence, to match with the solution far ahead of the wave, supposing that we take *z* = 0 to be the position of the front, we have the matching condition:

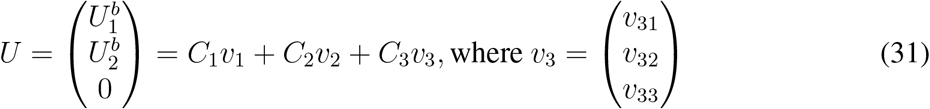

We obtain, 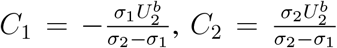 and *C*_*3*_ = 0 (details shown in Appendix C). Hence, we hypothesize that the reason why we obtain constant profile constant speed traveling waves is that, to match with the solution behind the wave, solution trajectories fall into the stable plane of the solution ahead of the front.

### 4.2. Comparison of wave speed generated from numerical simulations of the full system with the analytical solution derived from the linearized system

This section examines differences in wave-speed profiles calculated numerically from solving the full nonlinear system, with our analytically derived wave-speed, across various combinations of *α* and *β*, where *α, β* = 0, 1, …, 15 in the ECM dependent phenotypic switching functions *α*(*e*) = *αe* and *β*(*e*) = *β*(1 − *e*). Numerical solutions of the nonlinear system with zero degradation rate (*λ* = 0) indicate that, for small phenotypic switching rates, the wave speed is asymmetric (Figure 2a), which is inconsistent with the expression for 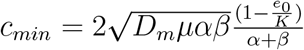 derived above. However, symmetry in wave speed emerges for higher values of *α* and *β* above 5. The maximum wave speed occurs along *α* = *β*, consistent with the analytical wave speed expression 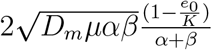. Notably, for a non-zero degradation rate (*λ* = 5), the speed does not reach its maximum at *α* = *β*. Instead, the maximum wave speed follows a line in the (*α, β*) coordinate system with a gradient less than 1 (Figure 2b). However, in fast phenotypic switching regimes, the traveling wave (TW) speed exhibits a similar pattern for the analytic expression and the numerically solved full nonlinear system (first and second panels of Figure 2c). It is interesting to note that for a large value of *α* and *β* the differences between the analytical solution and numerical solution of the TW speed are negligible (third panel of Figure 2c). Examination of the effect of degradation rate on TW speed in the fast phenotypic switching regime reveals no significant change between *λ* = 0, 5 and 10 for different choices of *α* and *β* (Figures 2d and f) in the fast phenotypic switching regime. We see that for large values of *α* and *β* (i.e., the fast phenotypic switching regime), the prediction of our linear analysis that the speed does not depend on *λ* is validated by our numerical results. However, if the timescale for matrix degradation is fast (that is, *λ* is large), then the percentage change of the TW speed increases for a certain regime of *α* and *β* (second panel of Figures 2d and f). This is what we would expect as an increase in the degradation rate creates additional space for migration (Figure 2e). However, we note that the difference between the numerically calculated wave speed and the analytic approximation is still relatively small.

**Figure 2:**
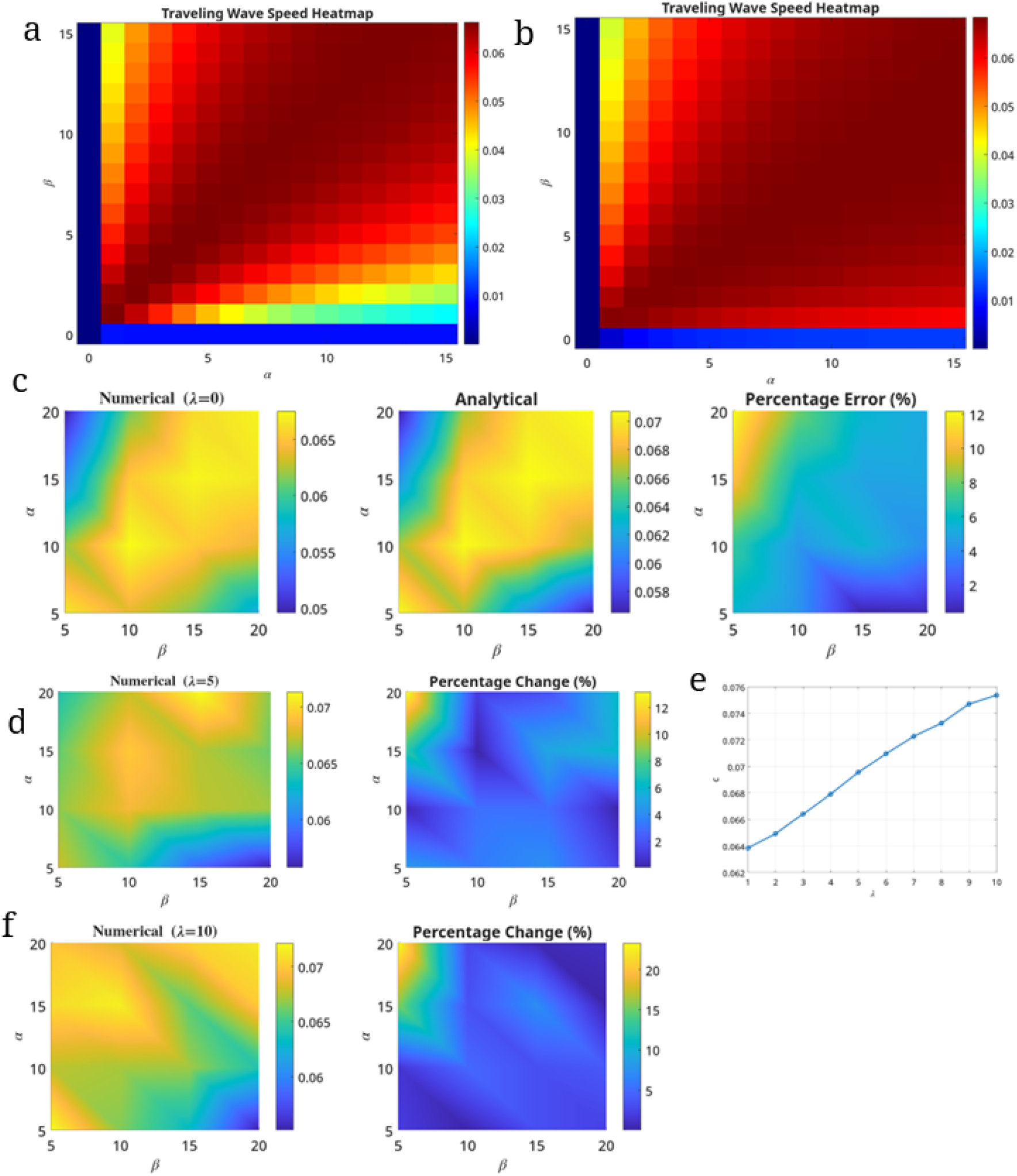
Heatmaps of wave speed for different values of *α* and *β* for (a) *λ* = 0 and (b) *λ* = 5. (c) The wave speed from numerical solution of Eqs (2)-(4) for *λ* = 0, the analytical expression of wave speed 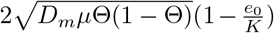, and absolute error between the analytical and numerical estimates of the wave speed. (d) Heatmap for wave speed for *λ* = 5 and the absolute change between wave speeds for *λ* = 0 and *λ* = 5. The percentage change is measured by 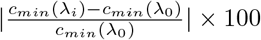 for *i* = 1, 2 with *λ*_0_ = 0, *λ*_1_ = 5 and *λ*_2_ = 10. (e) The effects of *λ* on wave speed for a fixed choice of *α* and *β*. (f) Heatmap for wave speed for *λ* = 10 and the absolute change between wave speeds for *λ* = 0 and *λ* = 10. Here, *D*_*m*_ = 0.02, *K* = 1 and *e*_0_ = 0.5 and *α*(*e*) = *αe* and *β*(*e*) = *β*(1 − *e*).

In addition, we examines the sensitivity of the TW speed to the values of migratory cell diffusivity and proliferative cell proliferation rate to the traveling wave (TW) speed within the full nonlinear model in the fast phenotypic switching regime. Linear stability analysis indicates that the minimal wave speed of the reduced model is proportional to the square root of *D*_*m*_*µ*, a relationship confirmed by numerical results. Furthermore, the speed of the full nonlinear model for *λ* = 0 closely matches the TW speed predicted by linear stability analysis (Figure S2a). For non-zero values of *λ*, analysis of the TW speed for various values of *D*_*m*_ and *µ* reveals no significant effects on TW speed, and the impact remains similar to that observed in the linearized system (Figure S2b).

## 5. Discussion

A tumor comprises multiple cell types with distinct phenotypes, each fulfilling specific roles within the tumor ecosystem, including proliferation and migration. Phenotypic switching among these cells is regulated by various tumor microenvironmental factors, such as oxygen tension, acidity, total tumor cell density, and local extracellular matrix (ECM) density [3, 8, 18, 24]. A go-or-grow model for tumor invasion with ECM-dependent phenotypic switching is proposed. In this model, local ECM is assumed to impede the diffusion of migratory cells. The go-or-grow model is formulated as a system of coupled partial differential equations (PDEs) for proliferative, migratory, and ECM densities. The possibility of reducing the go-or-grow model to a reduced model that eliminates the need for separate equations for migratory and proliferative cell types is investigated without altering the model’s essential dynamics. A relationship between the go-or-grow model and the reduced model is established in fast-phenotypic-switching regimes. Numerical results confirm that the reduced model accurately captures the dynamics of the go-or-grow model in these regimes for various phenotypic switching functions. Notably, the tumor composition varies depending on the choice of phenotypic switching. When both phenotypic switches (from proliferative to migratory and from migratory to proliferative) are constant functions, or only the proliferative-to-migratory switch is constant, both proliferative and migratory cells are present throughout the tumor. On the other hand, when only the migratory-to-proliferative switch is ECM-dependent, migratory cells are localized at the tumor front. Similar observations have been reported in the experimental literature [25, 26, 27] in the context of epithelial-to-mesenchymal transition (EMT), indicating that hybrid epithelial/mesenchymal state - that often overlaps with a basal breast cancer program - exhibit a more invasive phenotype. In addition, we observe that tumor invasion speed is high for ECM independent phenotypic switching from proliferative to migratory cell, but it is low for ECM independent phenotypic switching from migratory to proliferative cell type.

Irrespective of the phenotypic switching functions, numerical results suggest that the tumor propagates in a constant profile, constant speed traveling-wave manner. We carry out a traveling-wave (TW) analysis of the proposed go-or-grow model. Using linear stability analysis, we derived the minimal wave speed for the go-or-grow model, which is independent of the ECM degradation rate, and equal to the minimal wave speed obtained from the reduced model in the fast-phenotypic-switching regime. However, we noted, unlike the standard Fisher-KPP analysis, the steady state far ahead of the wave is unstable, leading to the hypothesis that the solution far behind the front forces the solution trajectories into the stable manifold region of the solution far ahead of the front. We have investigated numerically how the TW speed of the nonlinear system differs from the analytical solution. Our numerical results demonstrate that the difference between the TW speed for the nonlinear system and the linearized system is negligible when the ECM degradation rate *λ* = 0 in the fast phenotypic switching regime. More generally, we find that for non-zero *λ* and slow phenotypic switching, the linearized system still yields a good approximation for the TW speed of the non-linear system. Also, it is observed that the maximum value of the TW speed occurs when the rates of phenotypic switching are equal (i.e., *α* = *β*) for *λ* = 0. However, the maximum TW speed lies along a line, which makes a slope less than 45^°^ with the *α*-axis within the (*α, β*) phenotypic switching parameter space for *λ* = 5. Therefore, the maximum TW speed is attained for a relatively fast timescale of proliferative to migratory phenotypic switching than the timescale of migratory to proliferative phenotypic switching. In addition, we have noticed that as *λ* increases, the speed of the TW increases as one would expect intuitively. Moreover, we have explored how other parameters, such as diffusivity and proliferation rate, are sensitive to the wave speed of the nonlinear system in a way that agrees with our linear analysis.

Our proposed model suffers from various limitations. Firstly, it considers that the phenotypic switching in cells depends only on the local ECM density, and that there is no feedback on the properties of the ECM, other than density-degradation. Considering such feedback loops among the cell-matrix dynamics is an important next step [28]. Secondly, we do not consider the role of cellular memory, which has been observed for scenarios related to change in matrix stiffness, and also in the content of EMT where due to epigenetic modification, mesenchymal cells (migratory trait) maintain their phenotype even when they are in an environment favorable to switch to an epithelial phenotype (proliferative trait) [22, 29, 30]. Thirdly, we do not consider non-cell-autonomous dynamics such as co-operativity in migration that can drive invasion [31, 32, 33, 34]. Future work could examine how these factors influence tumor invasion dynamics and the traveling wave speed under ECM-dependent phenotypic switching.

## Data availability

The code for numerical simulations is available at GitHub link: https://github.com/gsadhuIITG/ECM-depends-go-or-grow.

## Acknowledgment

Param Hansa Philanthropies supports GS and MKJ. PKM would like to thank the Royal Society Yusuf Hamied Visiting Fellowship scheme, which funded his visit to IISc Bangalore.

## Appendix A. Numerical method

The domain [0, *L*] is divided into *N* equal grid spaces Δ*x*, with 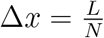, and Δ*t* is the time-step. We define 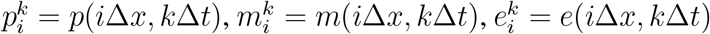 and 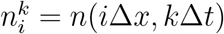. We employ forward difference scheme for the temporal derivative and central difference for the spatial derivative to discretize the governing equations.

The discretized form of Eq. (2) becomes

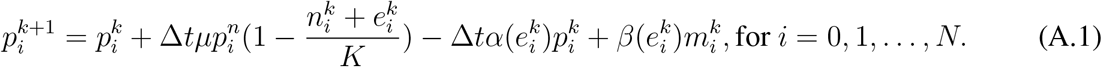

The discretized form of Eq. (3) becomes

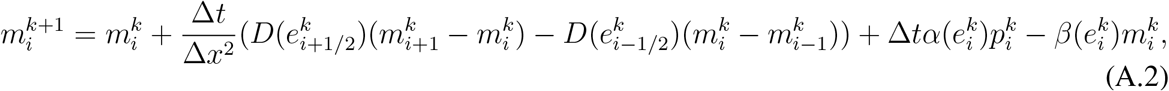

for *i* = 1, 2, …, *N* − 1. Here 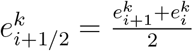 and 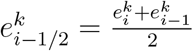.

At boundary points i.e., *i* = 0, *N*, we use ghost point criteria and the corresponding discretized form is prescribed as

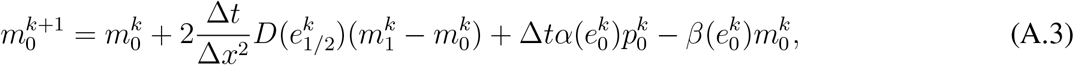

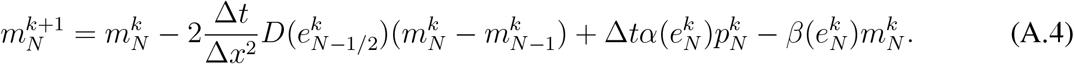

The discretized form of Eq. (4) is given as

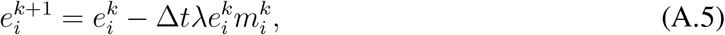

Numerical stability is ensured through the choice of time step length 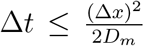 where *D*_*m*_ = max_*i*_ {*D*(*e*_*i*_)} in each time step.

The discretized form of Eq. (12) becomes

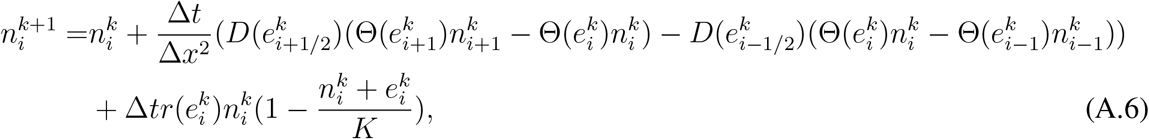

for *i* = 1, 2, …, *N* − 1. At the boundary points *i* = 0 and *N*, it takes the form

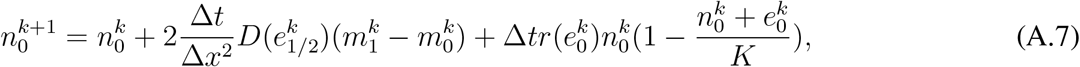

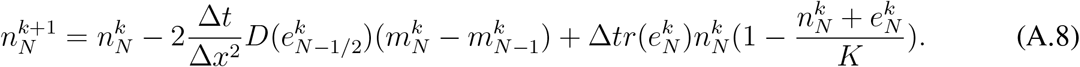

Eq. (13) is discretized as

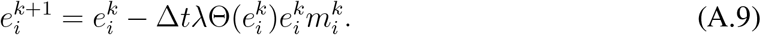

## Appendix B

Eq. (27) can be rewritten as

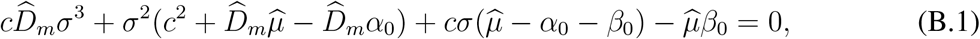

The condition for all roots of Eq. (B.1) to be real is

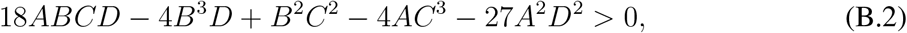

where 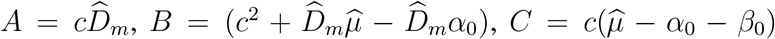 and 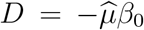. In the fast phenotypic switching regime, i.e., for large values of *α* and *β, B* and *C* become 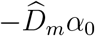 and −*c*(*α*_0_ + *β*_0_), respectively. Hence, the discriminant Eq. (B.2) becomes

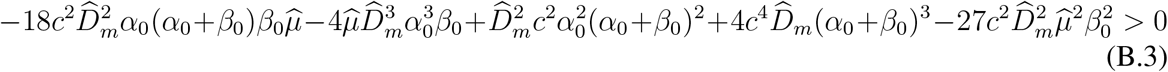

Dominant balance yields the condition

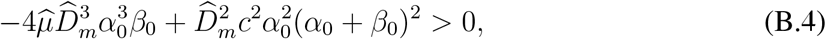

implying

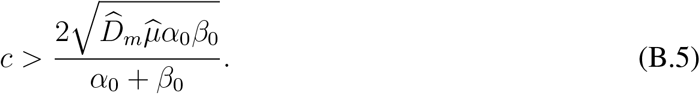

## Appendix C

From Eq. (31), we have

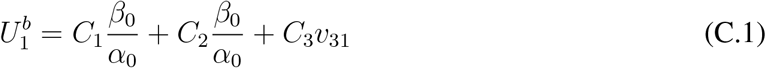

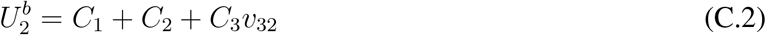

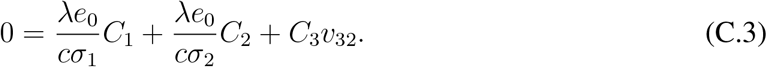

Since far behind the wave 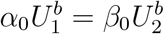 and from (C.1) and (C.2), we obtain,

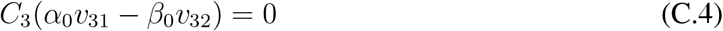

Hence, (C.4) gives either *C*_3_ = 0 or 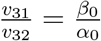.

Since,

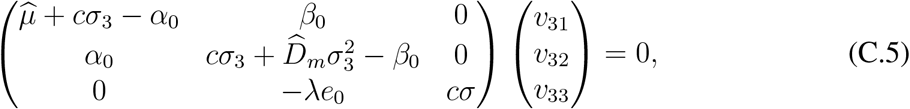

we have, 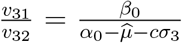, which obeys 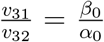 only when 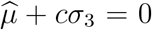 i.e., 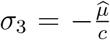. Since, 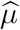 and *c* are both positive, this implies *σ*_3_ *<* 0, which is a contradiction as by definition.

Hence, (*α*_0_*v*_31_ − *β*_0_*v*_32_) ≠ 0. Therefore *C*_3_ = 0. From (C.1), (C.2) and (C.3), we obtain

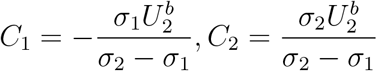

## Supplementary Information

**Figure S1:**
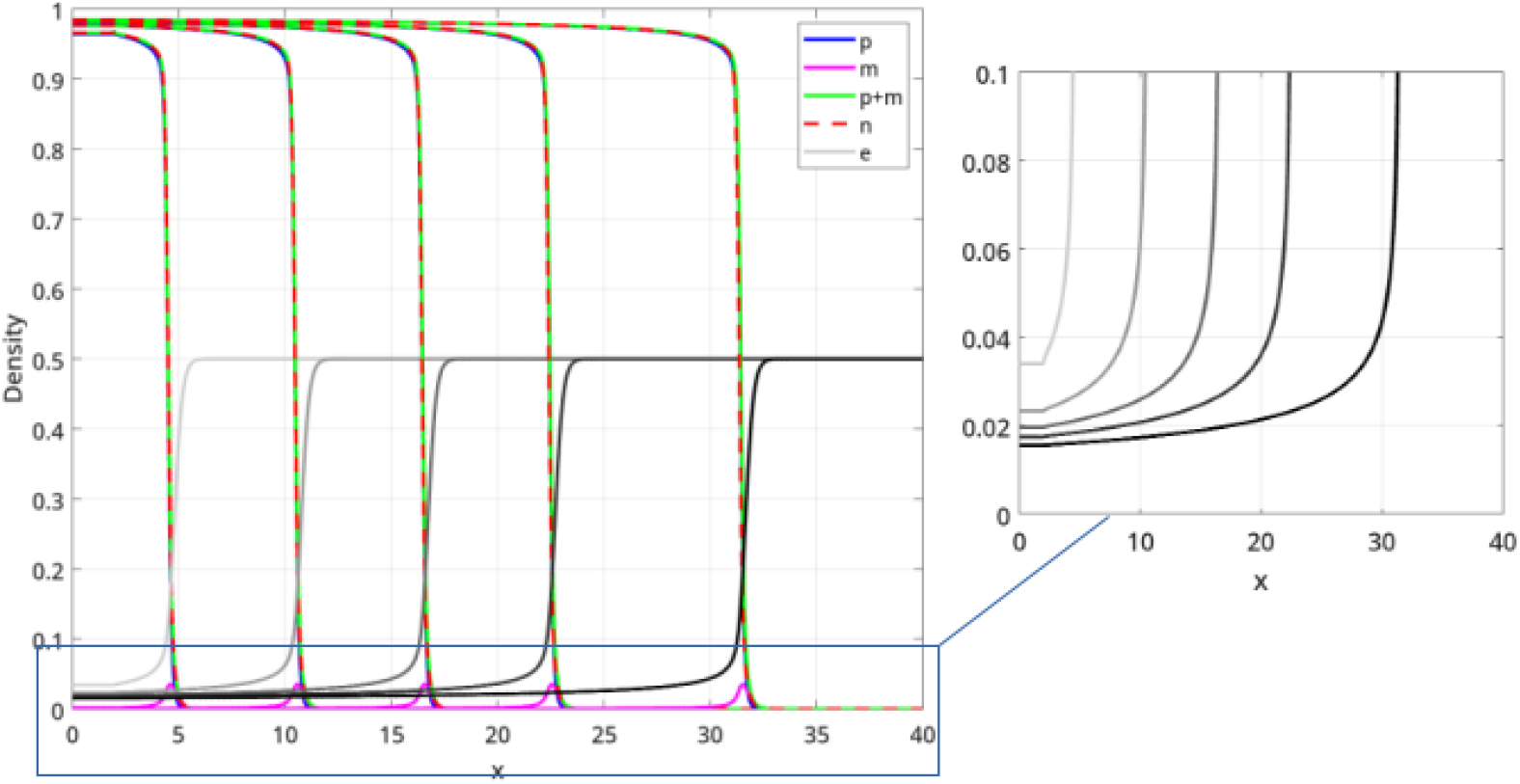
The spatial distribution of *p, m, p* + *m, n* and *e* at *t* = 50, 150, 250, 350 and 500. In the zoomed portion, ECM density distribution over the spatial domain at *t* = 50, 150, 250, 350 and 500.

**Figure S2:**
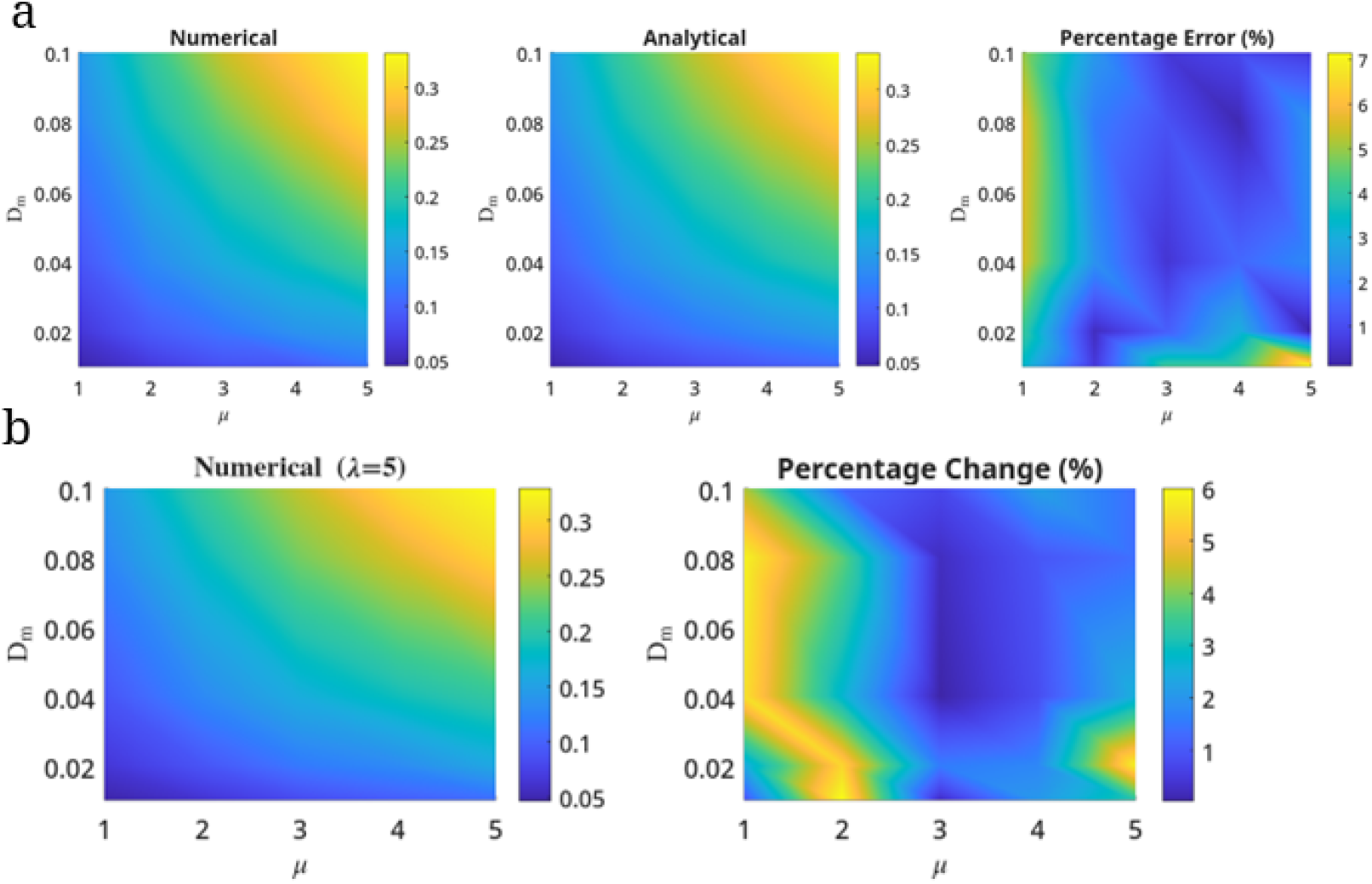
For different value of *D*_*m*_ and *µ*, (a) the wave speed from numerical solution of Eqs (2)-(4) for *λ* = 0, the analytical expression of wave speed 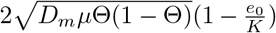, and absolute error between analytical and numerical estimate of wave speed. (b) Heatmap for wave speed for *λ* = 5 and the absolute change between wave speeds for *λ* = 0 and *λ* = 5. Here *K* = 1 and *e*_0_ = 0.5 and *α*(*e*) = 10*e* and *β*(*e*) = 20(1 − *e*).

